# VIProDesign: Viral Protein Panel Design for Highly Variable Viruses to Evaluate Immune Responses and Identify Broadly Neutralizing Antibodies

**DOI:** 10.1101/2025.05.21.654924

**Authors:** Tatsiana Bylund, Mohammed El Anbari, Andrew J Schaub, Tongqing Zhou, Reda Rawi

## Abstract

**Motivation:** Highly mutable viruses continuously evolve, with some posing major pandemic risks. However, standardized neutralization assays and up-to-date viral panels are often lacking, limiting evaluation of immunogens and identification of broadly neutralizing antibodies. Closing these gaps is essential for guiding effective countermeasure development.

**Results:** In this study, we present Viral Protein Panel Design (VIProDesign), a computational tool for designing viral protein panels that address the high sequence diversity of rapidly evolving viruses. VIProDesign uses the Partitioning Around Medoids (PAM) algorithm to select representative strains and applies the elbow-point method based on cumulative Shannon entropy to balance diversity and panel size. We used VIProDesign to generate optimized panels for Betacoronavirus, human immunodeficiency virus-1 (HIV-1), Influenza virus, Norovirus, and Lassa virus. The tool also supports customizable panel sizes, making it suitable for both resource-limited contexts and early-stage research. This flexible approach streamlines viral panel design across diverse pathogens. Although VIProDesign was originally developed for viral proteins, its underlying framework is broadly applicable to the selection of representative protein panels across diverse taxa, including bacterial species, toxins, and other biologically relevant protein families.

## 1 Introduction

Many viruses with pandemic potential, including Betacoronaviruses, HIV-1, and Influenza, exhibit substantial genetic variability and continuous evolution (Altman, et al., 2019; Bowen, et al., 2000; Chhabra, et al., 2019; Forni, et al., 2020; Hemelaar, 2012; Markov, et al., 2023; Mugosa, et al., 2016; Sarker and Jahan, 2023; Singh and Yi, 2021; Xue and Bloom, 2020). However, standardized panels for evaluating broadly neutralizing antibodies (bNAbs) and immunogen responses remain largely unavailable for most of these viruses due to significant cost and time constraints. The use of diverse pseudovirus strains across different studies further hinders the comparability of experimental results across studies. For instance, a panel of 60 HIV-1 isolates (Brown, et al., 2005) included 10 strains from each of the following six clades: A, B, C, D, CRF01_AE, and CRF02_AG, selected primarily based on availability, without prior analysis of population-level sequence diversity. In contrast, our in-house panel of 208 HIV-1 pseudoviruses (Doria-Rose et al., 2012) encompasses strains sampled between 1983 and 2008, offering broader temporal and genetic representation. A global panel of twelve HIV-1 Env strains (deCamp, et al., 2014) was developed using lasso regression to capture serum neutralization observed in a larger panel of 219 viruses. Most recently, Mkhize et al. (2023) analyzed HIV-1 viruses from 218 clinical trial participants collected between 2016 and 2020, primarily representing clades B and C. Complementary studies by Wieczorek et al. (Wieczorek, et al., 2024; Wieczorek, et al., 2023) proposed two focused panels: one comprising 30 contemporary subtype B pseudoviruses, and another including 50 recent subtype G and 18 CRF02_AG pseudoviruses derived from 10 Nigerian individuals across three studies conducted between 2003 and the present. Improving the relevance and utility of viral panels requires the inclusion of both historical and current strains to capture the full breadth of genetic diversity. Similar challenges exist across other viral pathogens, where the absence of standardized, representative panels hinders cross-study comparisons (Arevalo et al., 2022; Boyoglu-Barnum et al., 2021; Leonard et al., 2024; Moin et al., 2022). Incorporating genetically diverse and temporally distributed strains would enable more accurate assessments of immune responses and facilitate the development of broadly effective interventions.

Selecting such representative sequence sub-sets often relies on clustering methods, which typically require users to define parameters such as sequence identity thresholds, the number of clusters (Balaban, et al., 2019; Edgar, 2010; Fu, et al., 2012; Mirdita, et al., 2021; Pipes and Nielsen, 2022; Wright, 2024), or other parameters (James, et al., 2018), which often necessitates prior estimation and iterative parameter optimization. ALFATClust (Chiu and Ong, 2022) dynamically determines an optimal cut-off threshold for each cluster by simultaneously evaluating cluster separation and intra-cluster sequence similarity. SpCLUST (Matar, et al., 2019) employs a combination of Laplacian eigenmaps for dimensionality reduction and Gaussian mixture modeling and uses BIC to define number of clusters.

In this study, we introduce Viral Protein Panel Design (VIProDesign), an in-silico tool for constructing viral panels that capture the genetic diversity of highly variable pathogens while balancing strain coverage and panel size. VIProDesign also enables the generation of panels with user-defined sizes, making it particularly suitable for resource-limited settings or early-phase studies focused on candidate bNAb screening. Using this framework, we designed optimized panels for Betacoronavirus, HIV-1, Influenza virus, Lassa virus, and Norovirus. To demonstrate its broader applicability beyond viral proteins, we applied VIProDesign to a dataset of three-finger toxins and successfully identified a representative panel. These results highlight VIProDesign as a scalable and reproducible framework for panel design, with broad utility across diverse applications, including vaccine development, antivenom generation, and therapeutic discovery.

## 2 Materials and methods

### 2.1 Data

All sequences analyzed in this study were obtained from the GenBank database between October 2022 and February 2025.

Specific search queries were constructed for each viral protein dataset:

#### Betacoronavirus

(‘Betacoronavirus’[Organism] OR betacoronavirus[All Fields]) AND spike[All Fields] AND glycoprotein[All Fields]

#### Influenza

(“Homo sapiens”[Organism] OR Human[All Fields]) AND Influenza[All Fields] AND HA[All Fields]

#### Norovirus

(“Norovirus”[Organism] OR norovirus[All Fields]) AND (“Homo sapiens”[Organism] OR human[All Fields]) AND VP1[All Fields] NOT partial[All Fields]

#### Lassa Virus

(“Mammarenavirus lassaense”[Organism] OR Mammarenavirus lassaense[All Fields]) AND glycoprotein[All Fields]

#### HIV-1

Sequences were sourced from the HIV-1 Sequence Compendium 2021, curated by Los Alamos National Laboratory (Linchangco et al., 2021).

#### Three-finger toxins

Data were retrieved from UniProt (https://www.uniprot.org) using the query: (“three finger toxin” OR “3FTx”) AND keyword:“Toxin [KW-0800]”, with sequences filtered by length (60–100 amino acids). All datasets used in this study are publicly available via Zenodo (https://doi.org/10.5281/zenodo.15389017).

### 2.2 Preprocessing

Sequences with non-standard amino acids including ‘X’ or partial sequences were excluded. Sequence redundancy was minimized using CD-HIT (Li and Godzik, 2006) at a 99% identity threshold. Outliers were detected and removed using the Density-Based Spatial Clustering of Applications with Noise (DBSCAN) algorithm. The optimal value of ε for DBSCAN for determining outliers was defined as the elbow of k-distance graph (Kumar, 2024). However, for datasets with limited sequences, outlier detection was omitted to preserve biologically relevant diversity.

### 2.3 Clustering

The preprocessed sequences were clustered over a specified range of k values using the Partitioning Around Medoids (PAM) algorithm (Kaufman, 1990), which selects a medoid, an actual data point that represents the central element of each cluster. Panels of sequences were constructed for each value of k, using the k medoids.

### 2.4 Entropy Analysis

Sequences within each constructed panel were aligned using the AlignSeqs function from the DECIPHER R package unless a custom alignment was provided. The Shannon entropy was calculated at each alignment position, and the cumulative sum of Shannon entropy across positions was computed to assess sequence diversity. The optimal panel size was identified using the elbow-point method, balancing maximum diversity with manageable panel size. Alternatively, the user can select a predefined panel size. The clustering_info reports the number of viruses in each cluster, along with key cluster statistics.

## 3 Results

The design of comprehensive panels using VIProDesign follows the following procedure (Fig. 1). First, a set of sequences representing the target protein is assembled. The input data is then pre-processed by removing those containing unknown or ambiguous amino acid, typically represented by ‘X’. Next, sequence redundancy is removed by applying CD-HIT (Li and Godzik, 2006) with a 99% cutoff. The user also has the option to remove the outliers using DBSCAN algorithm (Kumar, 2024). If a predefined number of clusters, *M*, is specified, the PAM clustering algorithm is applied with *M* clusters, and the final panel consists of M cluster medoids. Alternatively, if a maximum number of clusters, *k,* is set instead, PAM clustering is performed for each value from two to k, and the final panel is selected at the elbow point of cumulative Shannon entropy.

**Figure 1.**
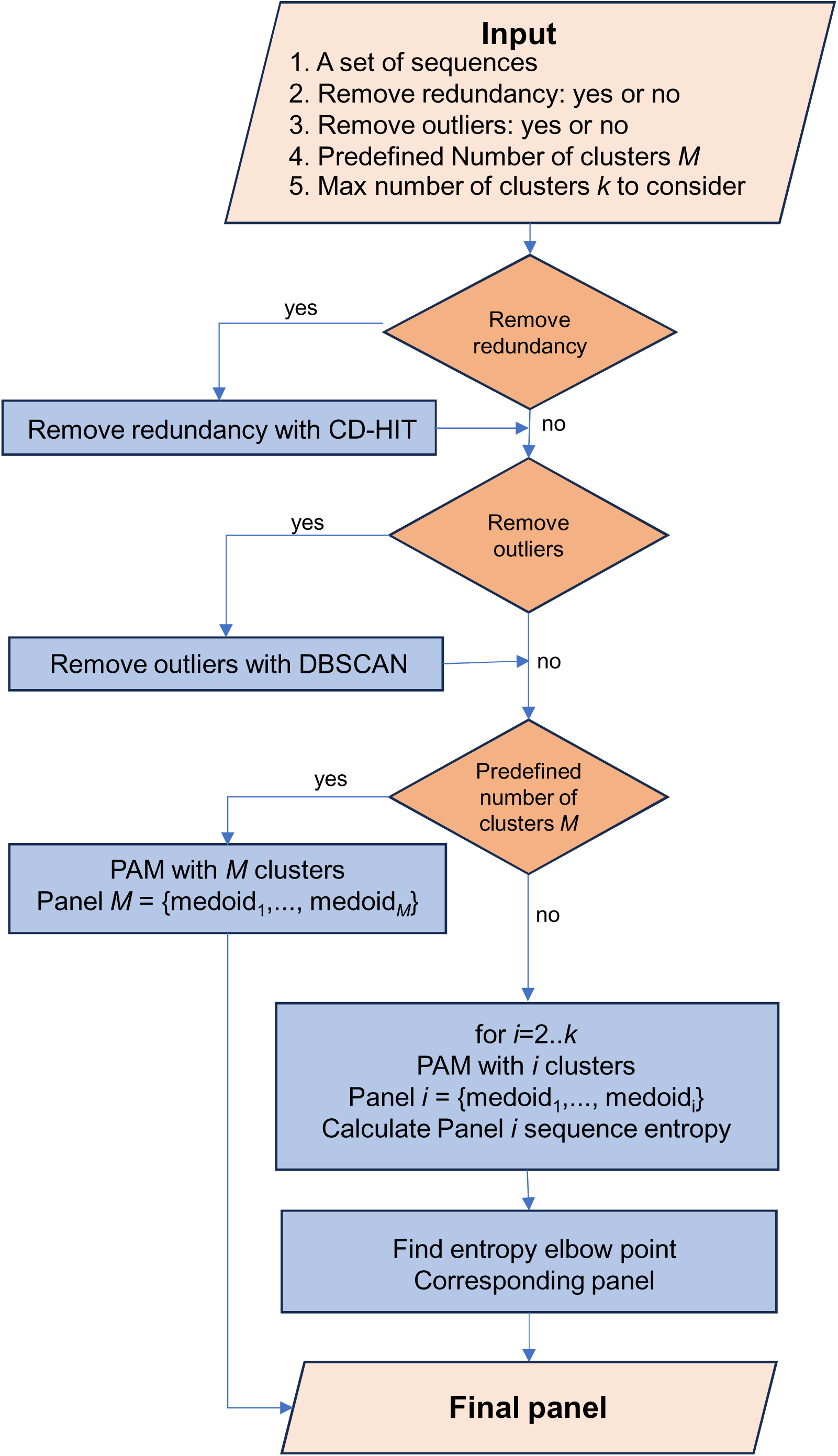
Flowchart of VIProDesign.

We applied VIProDesign to generate comprehensive panels for several highly variable viruses, including Betacoronavirus, HIV-1, Influenza virus, Norovirus, and Lassa virus. The proportion of outliers ranged from 3% for HIV-1 to 16% for Betacoronavirus (Supplementary Fig. 1). Due to the limited number of Betacoronavirus strains and higher proportion of outliers, we retained all Betacoronavirus sequences for the clustering step. Each viral dataset was clustered into k clusters using PAM, with *k* values ranging from 2 to 100, or up to 200 for HIV-1 due to its larger dataset size). A panel of *k* pseudoviruses was then designed, consisting of the *k* medoids from each cluster. The total Shannon entropy was calculated and plotted against the number of clusters for each *k* (Fig. 2). The optimal panel was selected at the elbow point of the entropy plot are named using the convention VIProDesign*_<VIRUS name>_<NUMBER of strains>*.

**Figure 2.**
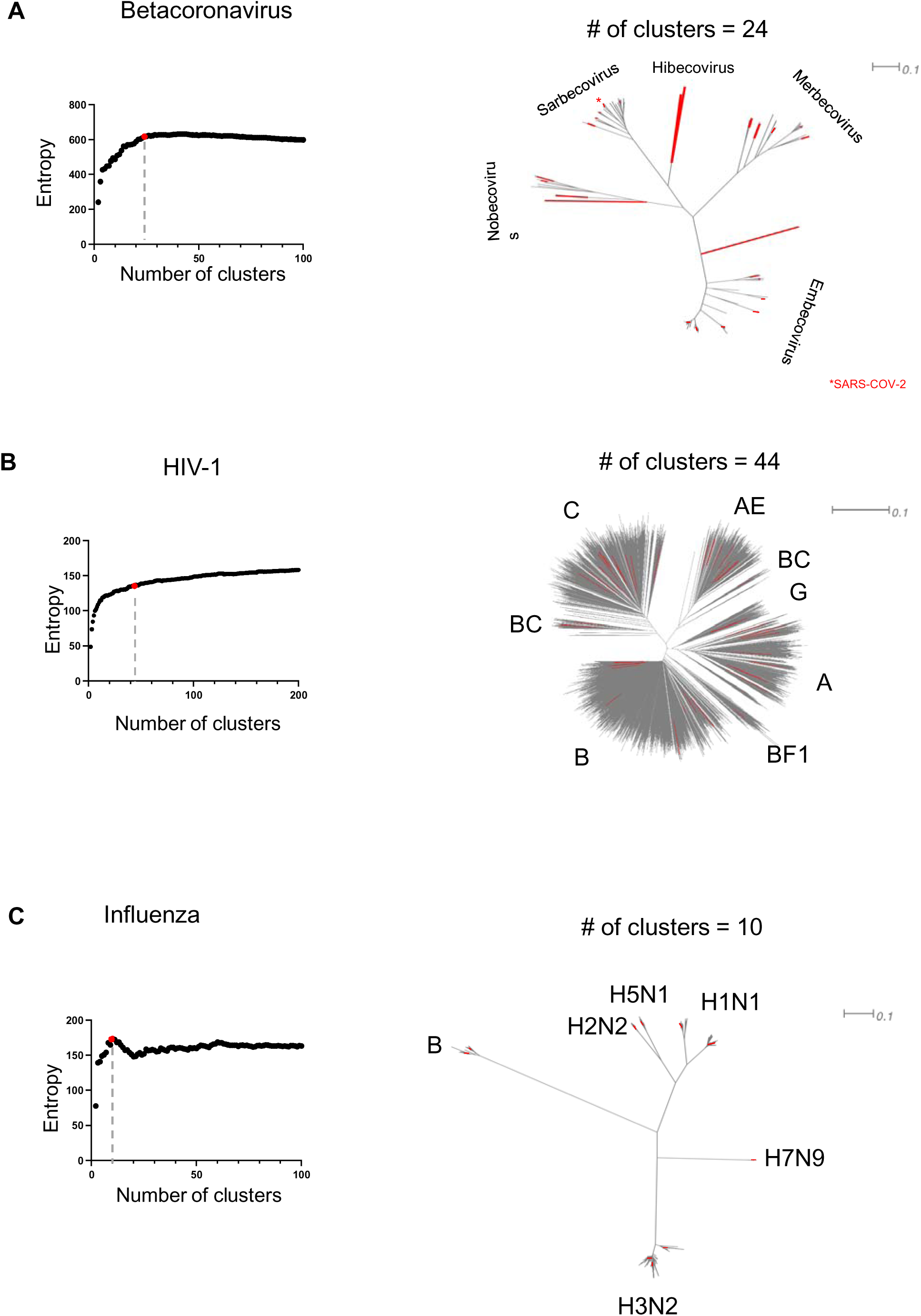
Panel designed for A) Betacoronavirus, B) HIV-1, C) Influenza. The sum of entropies is plotted against the number of clusters, with the elbow point highlighted in red in each graph. A phylogenetic tree representing the entire dataset is displayed, and the cluster centers corresponding to the elbow point are marked in red.

The VIProDesign_Betacoronavirus_24 panel comprised 24 strains (Fig. 2a), including five Sarbecovirus, four Nobecovirus, two Hibecovirus, five Merbecovirus, and eight Embecovirus strains. VIProDesign_HIV-1_44 panel consists of 44 strains (Fig. 2b), representing all major clades: A, AE, B, BC, BF1, C, G, and D.

VIProDesign_Influenza_10 panel includes 10 strains: two H1N1 and three H3N2 strains, one H5N1, one H2N2, one H7N9, and one Influenza B strain. Similarly, comprehensive panels were designed for Norovirus (18 strains) and Lassa virus (10 strains) (Supplementary Figure 2, Table 1).

Our in-house HIV-1 208-pseudo virus panel for assessing bNAbs breadth and potency is often impractical during early-stage discovery, when hundreds of antibodies must be screened. To address this limitation, we used VIProDesign pipeline to derive a reduced panel from the 208-pseudovirus panel, resulting in VIProDesign_HIV-1_18 panel (Figure 3). The panel includes strains from all major clades – A, AG, AE, B, BC, C, D, and G. Despite its dramatically reduced size, VIProDesign_HIV-1_18 panel showed strong predictive power of antibody performance on the full 208-virus panel. Across 12 well-characterized bNAbs – PGT121, PGT151, PGT145, VRC01, CH235.12, CAP256V2LS, DH270_lb, HERH-c.01, 8ANC195, 35O22, 2F5, and 10E8 – VIProDesign_HIV-1_18 achieved high correlations with full-panel results (Fig. 3c, d), with R² values of 0.94 for breadth (defined as IC80 < 1 µg/mL) and 0.96 for potency. These antibodies span diverse lineages and epitope specificities, and similar performance was observed in an expanded analysis of 53 bNAbs (Supplementary Table 2, Fig. 3). To accommodate resource-constrained settings, we further derived the VIProDesign_HIV-1_9 panel. While predictive performance is modestly reduced (R² = 0.83 for breadth and 0.94 for potency), the 9-pseudovirus panel remains a practical and informative surrogate for full-panel assays (Supplementary Fig. 3).

**Figure 3.**
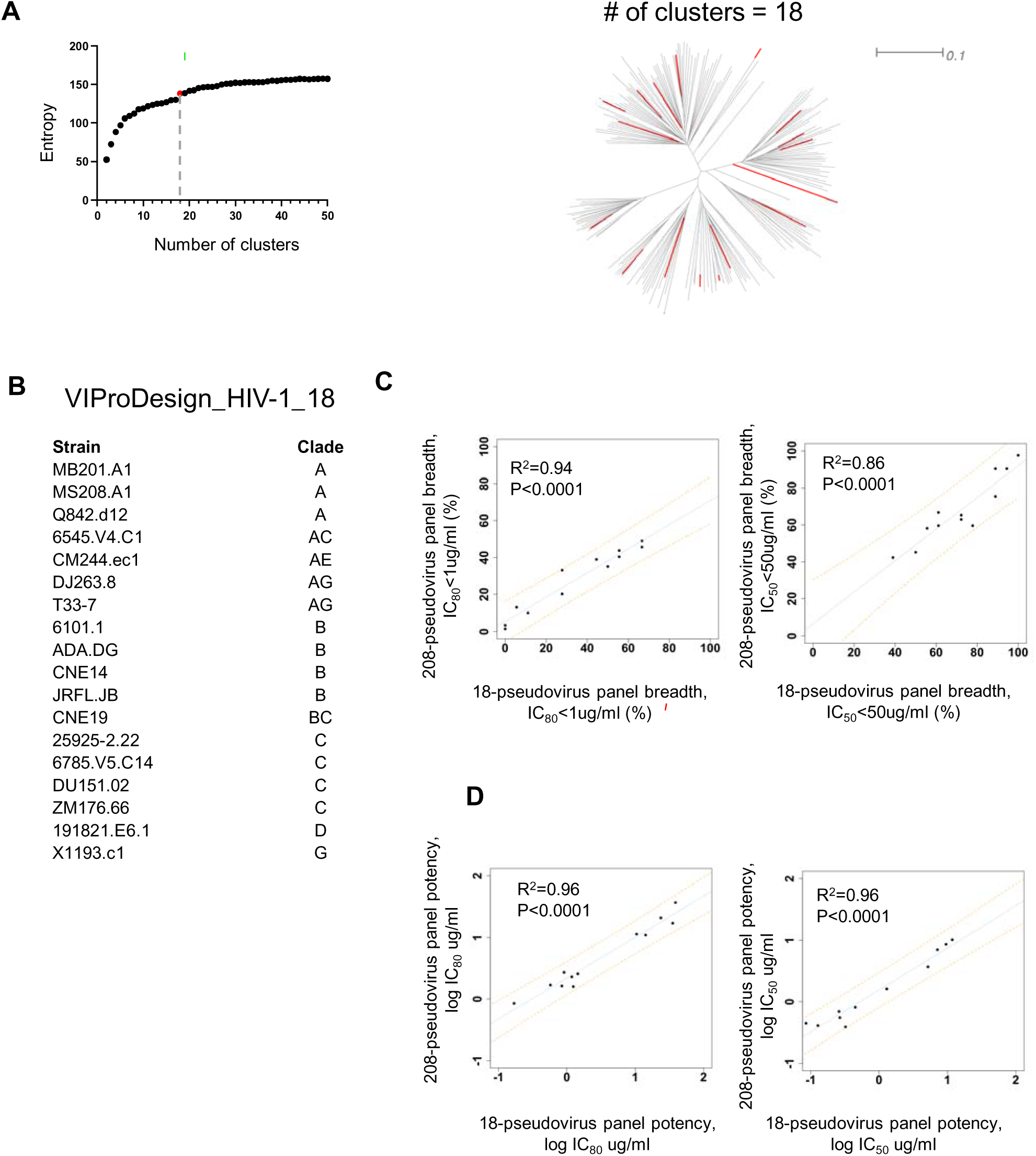
VIProDesign_HIV-1_18 panel designed based on commonly used HIV-1 208-pseudovirus set. A) The sum of entropies is plotted against the number of clusters. Elbow point = 18 is highlighted in red. The phylogenetic tree of 208 pseudoviruses is shown with 18 selected pseudoviruses highlighted in red. B) Designed HIV-1 panel. C) Linear regression of breadth (IC_80_<1 ug/ml and IC_50_<50 ug/ml) for 12 HIV-1 antibodies calculated based on 208-pseudovirus panel vs 18-pseudovirus panel. Predictive intervals are represented by orange dotted lines. D) Linear regression of potency for 12 independent HIV-1 antibodies calculated based on 208-pseudovirus panel vs 18-pseudovirus panel.

Given that some existing HIV-1 panels focus exclusively on individual clades (Hraber et al., 2017; Wieczorek et al., 2023, 2024), we applied VIProDesign specifically to HIV-1 clades B and C (Supplementary Fig. 4). The resulting clade-specific panels are temporally and geographically diverse. The clade B panel VIProDesign_HIV-1_B_40 includes strains sampled between 1981 and 2018, while clade C panel VIProDesign_HIV-1_C_65 spans isolates collected from 1989 to 2018, with both panels incorporating strains from multiple countries.

Encouraged by the results obtained for the HIV-1 panels, we sought to generate a small Influenza panel that could accurately recapitulate antibody breadth and potency as measured against our in-house 55-virus panel (Creanga, et al., 2021). Applying VIProDesign on the 55-virus panel initially yielded a set of eight pseudoviruses, comprising two H1N1 and two H3N2 strains, one H2N2, one H5N1, one H7N9, and one H10N8. Given the biased distribution of H1N1 (n=23) and H3N2 (n=26) strains in the full panel, we subsequently applied the panel separately to H1N1 and H3N2 to ensure a representation of their diversity. This yielded six H1N1 and six H3N2 strains. We substituted the initial two H1N1 and two H3N2 with six H1N1 and six H3N2 strains, resulting in a final 16-virus. The optimized panel demonstrated strong predictive performance, achieving R^2^ values of 0.83 for breadth (IC_80_ < 1 µg/ml) and 0.94 for potency across 24 antibodies from the reference study (Supplementary Figure 4).

To further test the generalizability of VIProDesign beyond viruses, we applied it to a curated dataset of three-finger toxins, resulting in a representative panel of 20 toxin sequences (Figure 5). This demonstrates the framework’s potential utility for designing protein panels in toxinology and other non-viral applications.

**Figure 4.**
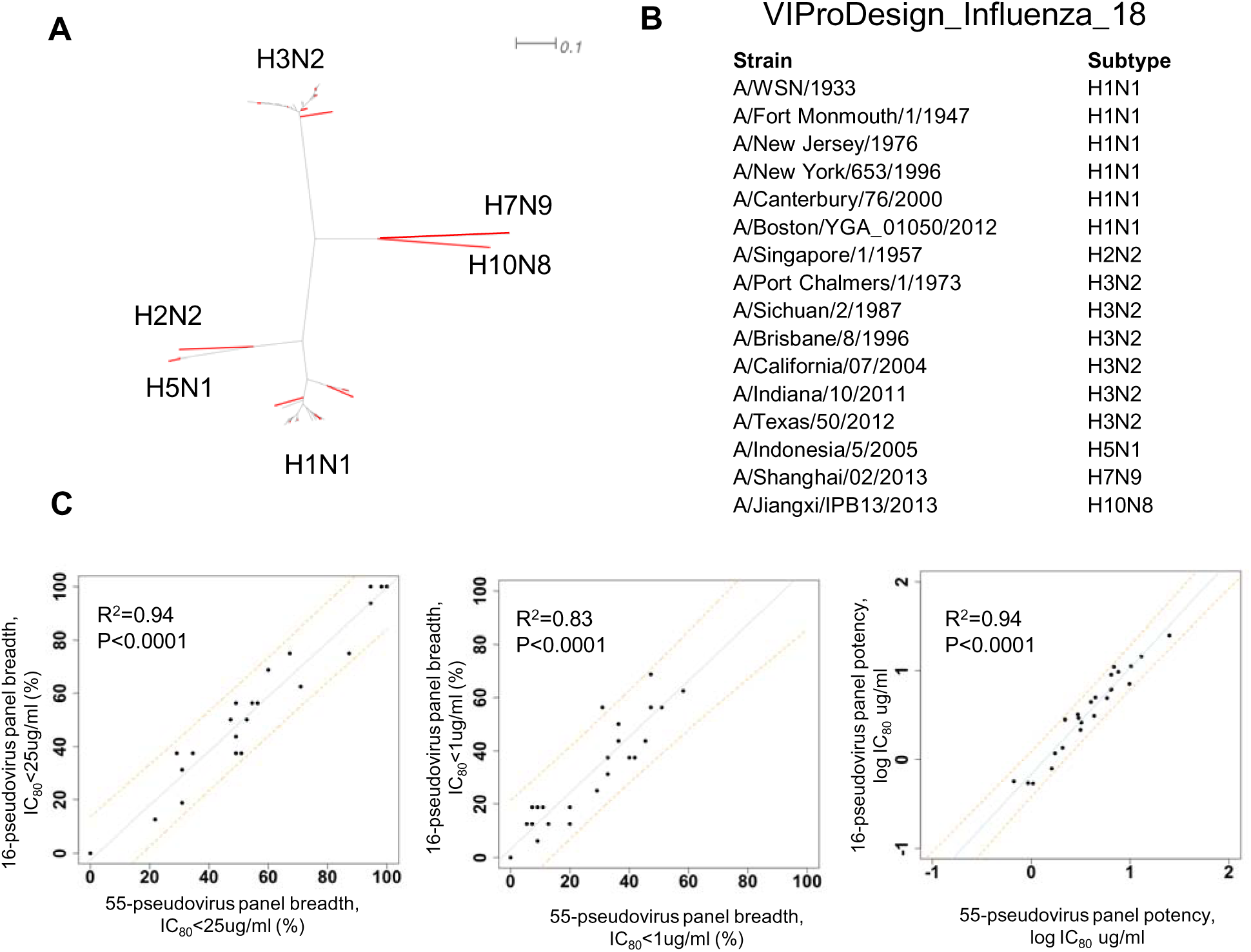
Influenza HA panel designed based on 55 Influenza panel. A) The phylogenetic tree of 55 strains is shown with 16 selected strains highlighted in red. B) Designed Influenza panel. C) Correlation of breadth (IC_80_<25 ug/m, IC_80_<1 ug/ml) and potency for 24 antibodies calculated based on 55-pseudovirus panel vs 16-pseudovirus panel. Predictive intervals are represented by orange dotted lines.

**Figure 5.**
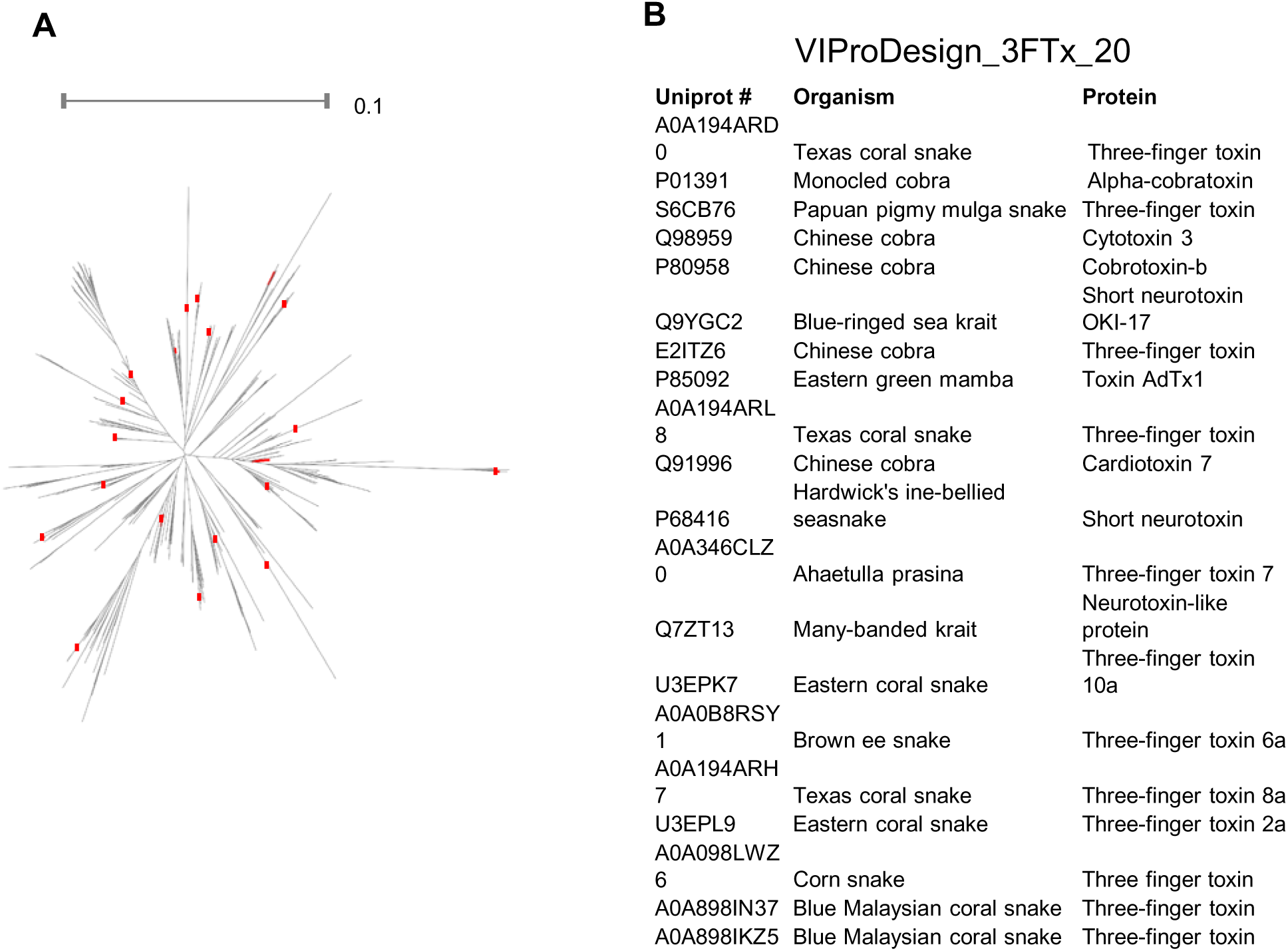
Three finger toxin panel. A) The phylogenetic tree of 560 three finger toxin sequences is shown with 20 selected strains highlighted in red. B) VIProDesign_3FTx_20 designed panel.

## 4 Discussion

Many viruses, including Betacoronavirus, HIV-1, and Influenza virus, remain significant public health threats due to their high variability and continuous evolution. When evaluating potential immunogens and broadly neutralizing antibodies, it is crucial to use a comprehensive virus panel that accurately reflects viral diversity and is highly representative. A common approach to capturing this diversity is clustering and selecting cluster representatives. However, many clustering algorithms require the number of clusters and other parameters to be specified. For example, ROBINEN clustering (Al Hasan, et al., 2009), which stands for ROBust INitialization based on inverse DENsity estimator, requires the number of clusters and MinPts (the number of nearest neighbors) as input parameters to identify dense regions during initialization. Since different viruses exhibit varying degrees of variation, selecting a single value for MinPts that works for all viruses is impractical. Our analysis has shown that ROBIN clustering is sensitive to outliers. Using DBSCAN for determining the number of clusters resulted in only four clusters for Lassa and HIV-1, which fails to capture their diversity. We also considered Affinity Propagation clustering (Frey and Dueck, 2007). Similar to DBSCAN, it identifies clusters based on density, but it determines the number of clusters in an unsupervised manner. Affinity Propagation clustering produced 167 clusters for the HIV-1 dataset, a number too large to be practical for experimental validation. In comparison, ALFATClust identified a single cluster for Lassa virus and 424 clusters for HIV-1. SpCLUST, on the other hand, identified one cluster for HIV-1, two clusters for Norovirus, and three clusters for Lassa.

PAM clustering algorithm used in this study selects a medoid to represent the cluster. Although this algorithm is less sensitive to outliers than k-means, our examination showed that it is still affected by outliers. Since the goal of this research is to design a panel of representative viruses, we aim to remove strains that are farther from others using DBSCAN.

PAM requires *n*, the number of clusters, as a parameter for the algorithm. To determine the final value of n, we performed clustering for a range of n = 2 to 100. A panel of size n consisted of n medoids, which were selected after performing PAM clustering with n clusters. To evaluate the diversity of a panel, we calculated sum of normalized Shannon’s entropy for the panel. Entropy increases rapidly at first (see Figure 2, Supplementary Figure 2), but then either slows down (HIV-1, Lassa) or decreases (Betacoronavirus, Influenza, Norovirus). An increase in entropy reflects an increase in diversity of the panel sequences. Slight differences in entropy values for clusters of 50 and above reflect the fact that the sequences no longer contribute significantly to increasing diversity. Since we aimed to cover the diversity of the virus while maintaining a reasonable panel size, we chose the elbow point of Shannon’s entropy as the final panel. Another benefit of using PAM clustering is that it allows for the design of a predefined panel of a specified size.

The designed VIProDesign Betacoronavirus panel consists of 24 strains from five Betacoronavirus subgroups: Nobecovirus, Sarbecovirus, Hibecovirus, Merbecovirus, and Embecovirus. Not all Betacoronavirus subgroups infect humans, although some animal-borne infections are known to present risk to humans (Chen, et al., 2025; Frutos, et al., 2021). When the DBSCAN outlier detection algorithm was applied to the Betacoronavirus dataset (Supplementary Figure 1a), the Nobecovirus and Hibecovirus subgroups were eliminated. This suggests that outlier detection is not suitable in this case, due to the limited number of strains (n = 200) in the Betacoronavirus set.

The designed VIProDesign Influenza panel consists of 10 strains. It has representatives from Influenza B, H1N1, H2N2, H3N2, H5N1 and H7N9. The different shape of the entropy graph (Figure 2c) with distinct pick reflects the fact that there is little variation within each subtype. The average sequence identity between Influenza HA is 34% ± 39%, while sequence identity between H3N2 sequences is 93% ± 8%.

Since the commonly used HIV-1 208-pseudovirus panel is impractical for many immunogen and antibody evaluation studies, we aimed to design a more compact panel that would predict antibody breadth and potency (Figure 3). The designed VIProDesign HIV-1 18 panel is highly predictive of antibody breadth and potency on 208-pseudovirus panel (Figure 3c,d, Supplementary Figure 3a). Some experiments may require even smaller number of pseudoviruses due to limitations in experimental resources. Since VIProDesign enables the design of panels with a predefined size, we designed a VIProDesign_HIV-1_9 panel derived from the 208-pseudovirus panel (Supplementary Figure 3b). This panel includes pseudoviruses from all major clades (A, AE, AG, B, BC, C, D, and G) and is highly predictive of antibody breadth and potency (Supplementary Figure 3c). All pseudoviruses, except for 217-11, which is highly similar to T33-7, are also present in the 18-pseudovirus panel, which is very useful for follow-up experiments. An intermediate-sized panel may also be useful, and more accurate.

The designed VIProDesign HIV-1 comprehensive panel consists of 44 strains from all major clades. The sequence identity between HIV-1 strains ranges from 58% to 90%, with an average identity of 75% ± 3% between sequences.

When examining panels of different sizes, we observed that smaller panels are often subsets of larger ones. This reflects the fact that PAM selects a medoid from each cluster, and thus the centers of densely populated area are chosen. This is extremely beneficial, if a smaller panel is used in the initial steps of the experiments, and a larger size panel – in the later steps. A smaller panel will consist of representatives from the most densely populated regions, thereby reflecting the greater part of virus diversity. VIProDesign also outputs the number of viruses in each cluster, which could be useful to consider when selecting a panel.

The specifics of virus subgroup variation may require separate considerations when designing a comprehensive panel. Whereas HIV-1, Norovirus, and Lassa exhibit more uniform variability, as seen in the phylogenetic trees (Supplementary Figure 1), Influenza viruses exhibit a higher sequence identity difference between subtypes, approximately 40%, and a lower difference within a subtype, around 10%. When designing a comprehensive panel based on 55-virus panel (Creanga, et al., 2021) we needed to consider H1N1 and H3N2 separately, as the panel contained a large portion of viruses from both subtypes. After including six pseudoviruses from both H1N1 and H3N2, the final 16-pseudovirus panel proved to be highly predictive of antibody breadth and potency (Figure 4c).

Clade B and C are two of the major clades of HIV-1. Clade B is most prevalent in North America, Europe and Australia (Wieczorek, et al., 2023). Clade C is the most common lineage (Hraber, et al., 2017). It is interesting to note that the designed comprehensive panels included strains from a range of years, spanning from 1981 to 2018 (Supplementary Figure 4). It has been shown that HIV-1 virus evolves to become more resistant to broadly neutralizing antibodies (Mkhize, et al., 2023). When designing comprehensive panels, it is important to consider all available strains, as viruses are known to be reintroduced (Carter and Sanford, 2012). For example, the sequence identity between A/Alany/4835/1948//HA (Genbank: ABN59401.1) and A/Kiev/59/1979//HA (Genbank: AAA43172.1) is 98%, despite a 31-year difference between them. The sequence identity between B.US.06.04013440_5_B11.GU330809 and B.US.16.B207_CMV_CD69_G13.MT190767 is 91%, despite a 10-year difference, whereas sequence identity between two HIV-1 Env sequences from Brazil from the same year (B.BR.10.10BR_PE020.KT427739 and B.BR.10.10BR_PE006.KT427745) is 78%.

In this work, we developed VIProDesign, a novel approach for designing comprehensive virus panels for highly variable viruses. Our method utilizes Partitioning Around Medoids to select representative viruses and determines the panel size using the elbow point of Shannon entropy to maximize the diversity of the panel, while maintaining a reasonable panel size. This approach also allows for the creation of panels with a predefined size, making it especially valuable when smaller panels are needed due to resource constraints.

While VIProDesign was developed with viral proteins in mind, our application to three-finger toxins highlights the framework’s broader utility for generating compact, representative panels for other biologically diverse protein families. Three-finger toxins are a structurally conserved but functionally diverse class of snake venom proteins, and panel optimization in this space could be valuable for antivenom discovery, structure-function studies, and immunological profiling. This use case demonstrates that VIProDesign is not limited to virology and can be readily applied to other domains, including toxinology, parasitology, and bacterial pathogenesis – any context in which highly diverse protein repertoires complicate comprehensive screening or therapeutic development.

## Acknowledgements

We thank J. Stuckey for assistance with the figures. We thank all members of SBS and SBIC VRC NIAID for their feedback. This work utilized the computational resources of the NIH HPC Biowulf cluster (NIH HPC). This study used the Office of Cyber Infrastructure and Computational Biology (OCICB) High Performance Computing (HPC) cluster at the National Institute of Allergy and Infectious Diseases (NIAID), Bethesda, MD.

## Funding

This work has been supported by the Intramural Research Program (National Institute of Allergy and Infectious Diseases, National Institutes of Health, USA).

## Conflict of Interest

*none declared*.

---

**Supplementary Figure 1.**
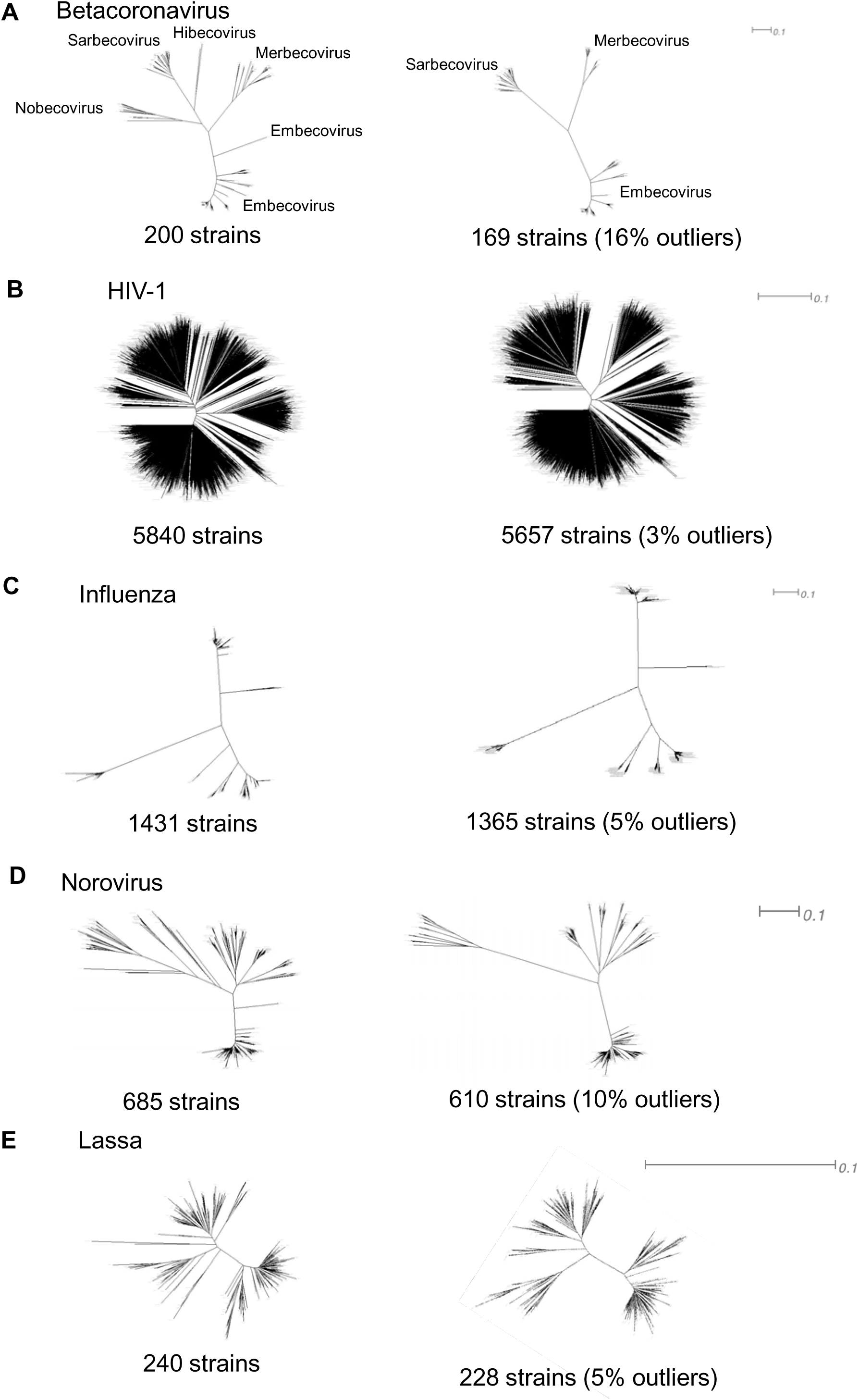
Detecting outliers with DBSCAN. Phylogenetic trees of complete dataset and dataset with outliers removed are shown for A) Betacoronavirus, B) HIV-1, C) Influenza, D) Norovirus and E) Lassa.

**Supp Figure 2.**
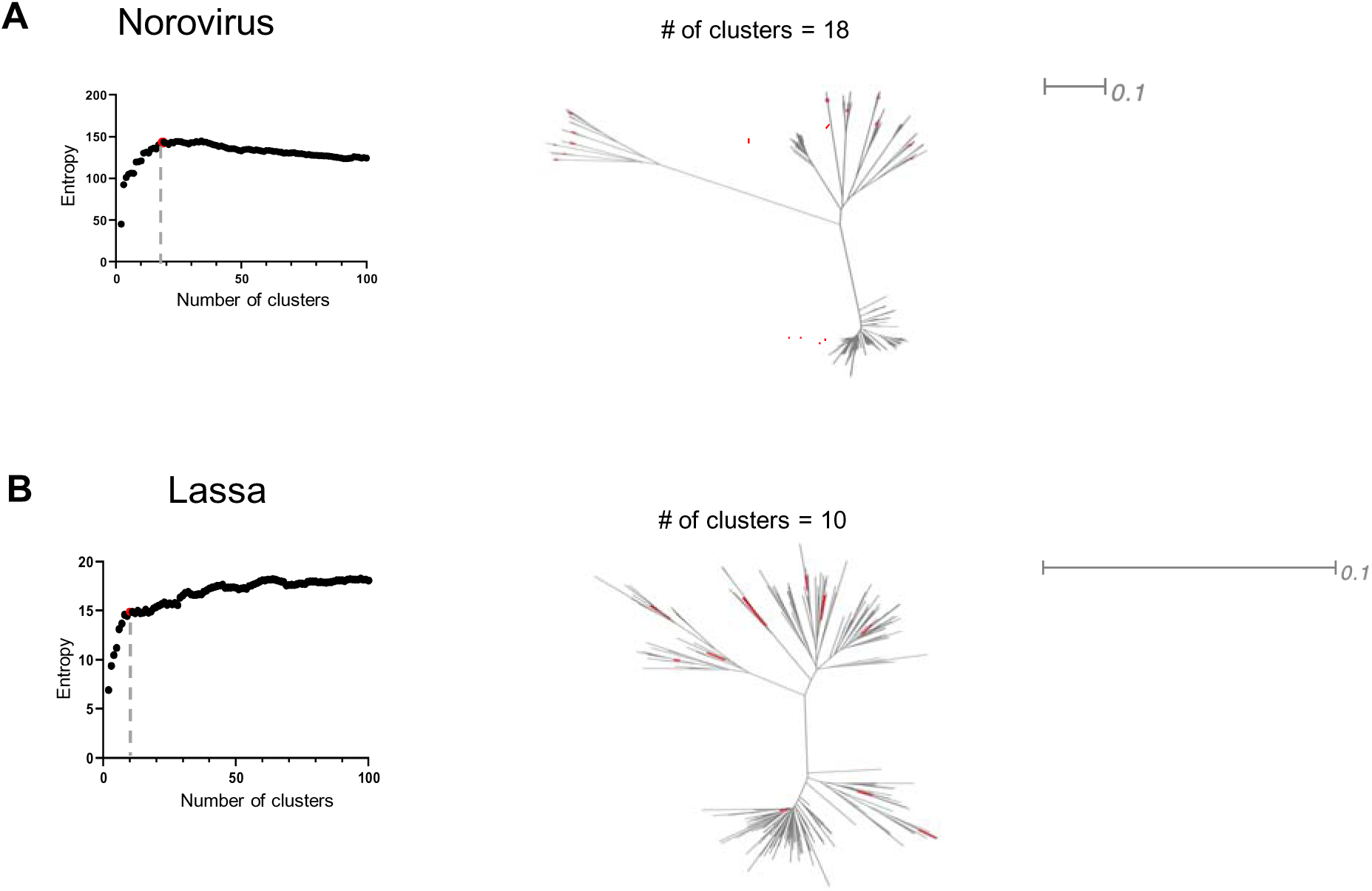
Panel designed for A) Norovirus, B) Lassa. The sum of entropies is plotted against the number of clusters, with the elbow point highlighted in red in each graph. A phylogenetic tree representing the entire dataset is displayed, and the cluster centers corresponding to the elbow point are marked in red.

**Supplientary Figure 3.**
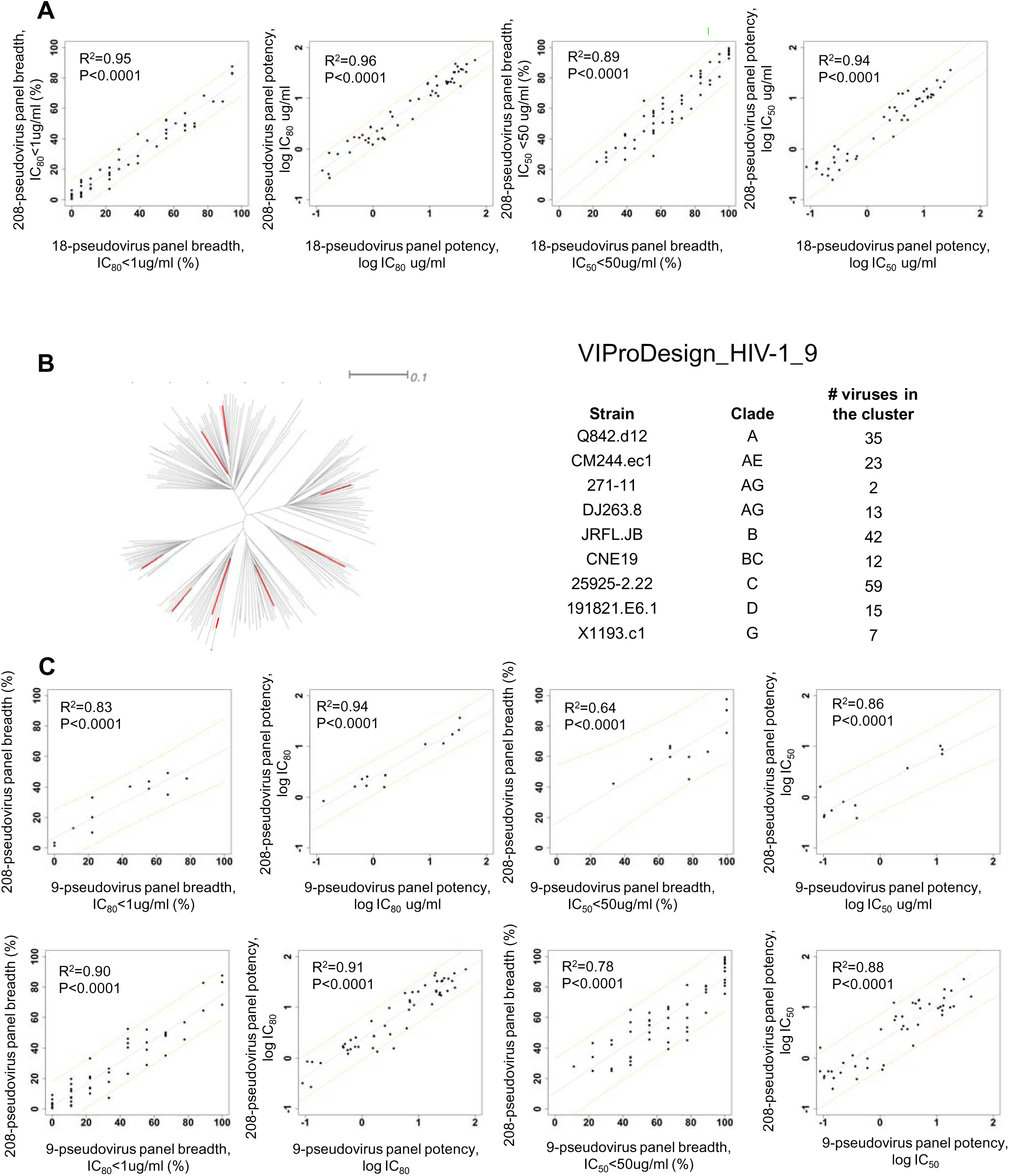
HIV-1 panel designed based on commonly used HIV-1 208-pseudovirus set. **A**) Correlation of breadth and potency (IC_50_<50 ug/ml) for 53 HIV-1 antibodies calculated based on 208-pseudovirus panel vs 18-pseudovirus panel. Predictive intervals are represented by orange dotted lines. **B**) The phylogenetic tree of 208 pseudoviruses is shown with 9 selected pseudoviruses highlighted in red. **C**) Correlation of breadth (IC_80_< 1 ug/ml and IC_50_<50 ug/ml) and potency for 12 independent HIV-1 antibodies and 53 HIV-1 antibodies calculated based on 208-pseudovirus panel vs 9-pseudovirus panel.

**Supplementary Figure 4.**
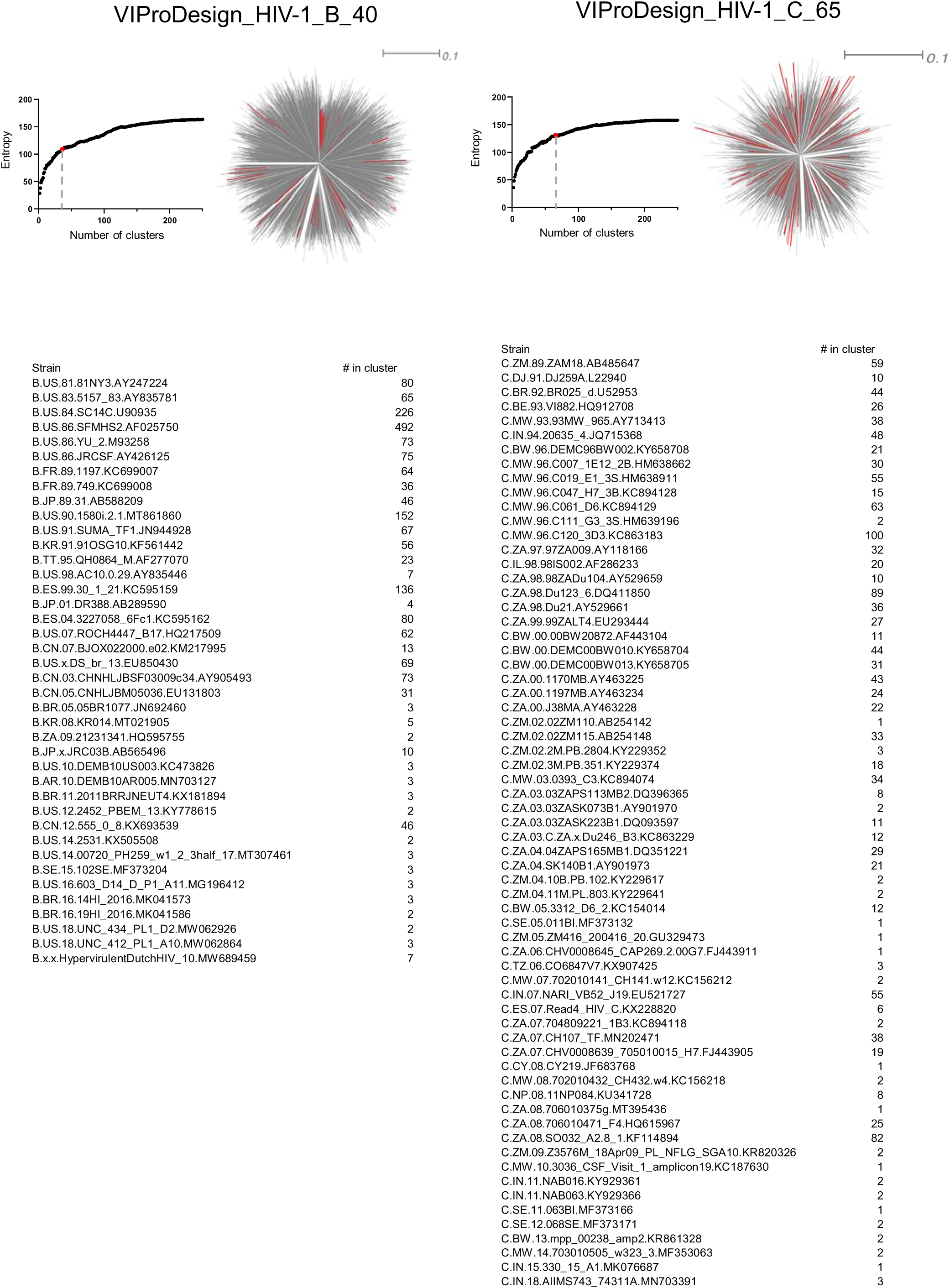
HIV-1 clade B and C panels.

## Notes

https://doi.org/10.5281/zenodo.15389017

### Competing Interest Statement

The authors have declared no competing interest.

### Summary of Updates

Some grammatical changes were made throughout the manuscript.

